# Absence of the fragile X mental retardation protein alters response patterns to sounds in the auditory midbrain

**DOI:** 10.1101/2022.01.15.475369

**Authors:** Jérémie Sibille, Jens Kremkow, Ursula Koch

## Abstract

Among the different autism spectrum disorders, Fragile X Syndrome (FXS) is the most common inherited cause of mental retardation. Sensory and especially auditory hypersensitivity is a key symptom in patients, which is well mimicked in the Fmr1 -/- mouse model. However, the physiological mechanisms underlying FXS’s acoustic hypersensitivity in particular remain poorly understood. Here, we categorized spike response patterns to pure tones of different frequencies and intensities from neurons in the Inferior Colliculus (IC), a central integrator in the ascending auditory pathway. Based on this categorization we analyzed differences in response patterns between IC neurons of WT and Fmr1 -/- mice. Our results report broadening of frequency tuning, an increased firing in response to monaural as well as binaural stimuli, an altered balance of excitation-inhibition, and reduced response latencies, all expected features of acoustic hypersensitivity. Furthermore, we were surprised to notice that all neuron response types in Fmr1 -/- mice displayed enhanced offset-rebound activity outside their excitatory frequency response area. These results provide evidence that the loss of *Fmr1* not only increases spike responses in IC neurons similar to auditory brainstem neurons, but also changes response patterns such as offset spiking. One can speculate this to be an underlying aspect of the receptive language problems associated with Fragile X syndrome.

## Introduction

After Down’s syndrome, Fragile X Syndrome (FXS) is a major cause for intellectual disability (Ciaccio et al. 2017), affecting approximately 1 in 4000 males and 1 in 6000 females (Turner et al. 1996). It originates from an over expansion of the CGG triplet repeats on the Fragile X mental retardation 1 (*Fmr1*) gene located on the X chromosome. This CGG over expansion leads to a loss of Fragile X Mental Retardation Protein (FMRP) resulting in a dysfunctional neuronal development. Patients with FXS display multiple abnormalities, including phenotypic features (e.g. protruding ears and macroorchidism), hypersensitivity to different sensory modalities, language retardation and social withdrawal (Finestack et al., 2009; Rais et al., 2018). A consensus has emerged that the animal model of Fragile X, the *Fmr1* -/- mouse, recapitulates most neurologic symptoms of FXS patients, with hypersensitivity to sounds and the development of audiogenic seizures as one of the key sensory dysfunctions (Dolan et al. 2013).

Audiogenic seizures in rodents originate in part from overactivity in the inferior colliculus (IC), which is a prominent midbrain nucleus and a central hub for sound processing. Lower auditory brainstem nuclei, including the cochlear nucleus (CN), the lateral superior olive (LSO), and the nuclei of the lateral lemniscus (LL) almost exclusively project to the central nucleus of the IC, which itself projects upward through the thalamus - the Medial Geniculate Nucleus – finally reaching to the Auditory Cortex (AC). Functionally, the IC mediates ascending streams of information, but also receives direct descending streams of cognitively-driven activity from corticofugal projections originating from layer V and VI of the AC and many non-auditory, cortical areas (Olthof et al., 2019; Yudintsev et al., 2021). These latter corticofugal projections likely support the plasticity observed in the IC when facing novelty in auditory stimuli (Ayala and Malmierca, 2013; Ayala et al., 2016). Such functions underpin the central role of the IC in many aspects of auditory-sensory processing, which may be directly affected in Fragile X related auditory dysfunctions. Indeed, a clear link has been proposed between audiogenic seizures during development and glutamatergic neurons in the IC by selectively deleting FMRP in VGluT2 neurons of the IC (Gonzalez et al., 2019).

Previous studies relate the known acoustic hypersensitivity with higher neuronal firing rates along the ascending auditory pathway of *Fmr1* -/- mice, as already shown in the LSO (Garcia-Pino et al., 2017), the inferior colliculus (Nguyen et al., 2020) and the auditory cortex (Rotschafer and Razak, 2013). The studies in the auditory brainstem strengthen the link between hyper-responsiveness of neurons and enhanced synaptic transmission at excitatory synapses together with reduced inhibitory input number and altered action potential properties caused by alterations in various potassium channels (Strumbos et al., 2010; Garcia-Pino et al., 2017; McCullagh et al., 2020). These physiological changes emerge after hearing onset and persist throughout adulthood (McCullagh et al., 2020).

The present study explores to what extent spike response patterns to pure tones across the entire frequency response area are altered by a deletion of *Fmr1*. Since changes in response strength and tuning bandwidth turned out smaller than expected, we categorized neurons based on their neuronal response patterns. We also explored whether binaural integration of sounds is affected by *Fmr1* deletion also at the midbrain and whether adaptational phenomena, such as stimulus specific adaptation, was impaired in *Fmr1* -/- mice.

## Materials and Methods

### Animal welfare

All experiments were performed with the approval of the German animal welfare legislation (Landesamt für Gesundheit und Soziales, LaGeSo Berlin T0262/12). Wild-type (WT) and *Fmr1* -/- mice colony founders were obtained from Jackson Laboratories (FVB.129P2-Pde6b_Tyrc-chFmr1tm1Cgr/J, stock #004624; RRID:IMSR_JAX:004624). Mouse lines were sustained in our animal facility, housed in a 12h/12h light/dark cycle, and had ad libitum access to food and drinking water. No conditioning or manipulations were performed prior to the recording day.

### Experimental design and statistical analysis

All efforts were made to minimize both the suffering and the number of used animals in this study. Overall, 37 mice (21 WT and 16 *Fmr1* -/-; littermates from both sexes) aged 52 to 120 days were used during this project. Statistical analyses were conducted with Matlab (RRID:SCR_001622). One-way or two-way ANOVA was applied when comparing age and genotypes while Post-hoc Bonferroni or Dunnett’s pairwise comparisons were performed only where two-way ANOVA gave statistically significant differences. The unpaired two-tailed Student’s t-test for normally distributed data was applied to probe differences between two means while Mann–Whitney test was used for pairwise comparison of non-parametrically distributed data.

### Surgery

Anesthesia was first induced by intraperitoneal injection of xylazine (Rompun 2 %, 5 mg/kg) and ketamine (Ketavet 10%, 100 mg/kg). Additional doses were given subsequently during the course of the experiment based on vibrissa motility and heart-beat frequency. Rectal probe helped to maintain the animal body temperature between 36° C and 37° C (TC01, Multi-Channel-System). The frontal skull was first exposed and cleaned. A locally custom-made head post was attached to the frontal part of the skull using dental cement (Charisma, Heraeus-Kulzer). The animal was then placed into a computer-controlled stereotaxic frame (Neurostar) in a sound-attenuating chamber (Desone) where the head-to-stereotactic-arm alignment was thoroughly adjusted every day. A craniotomy was performed caudally to lambda (0 to -2 mm Rostro-Caudal and -2 mm to 0 Medio-Lateral) exposing the cerebellum and the underlying IC. Great care was used during the craniotomy to follow the posterior border of the lateral suture in order to preserve the underlying blood vessel. Duratomy was performed, tissue wetted with PBS, and heartbeat was constantly monitored.

### Acoustic stimulation

Sound stimuli were designed using Brainware (TDT, Jan Schnupp). Sound stimuli were produced at the 195.3 kHz sampling rate of the RZ6 TDT amplifier and monitored by Real-Time Processor Visual Design Studio (RPvdsEx; Tucker Davis Technology TDT). Signals were delivered either monaurally or binaurally to the ears of the experimental animals by MF1 magnetic speakers (TDT, closed field configuration) and appropriate plastic tubing inserted into the animal’s pinnae. We calibrated our acoustic signals using a 1/4 inch measurement microphone (40 BF, G.R.A.S.), custom written Matlab software, and SigCalRP calibration software (TDT). Single neuron spiking activity was pre-amplified and band-pass filtered (300– 3000 Hz) using a Model 3000 amplifier (A-M Systems). Neuronal spikes were digitized at 24.4 kHz by the RZ6 Processor (TDT). Real-time spiking activity was gathered according to stimuli and displayed using Brainware. White noise (100 ms duration, 5 ms rise/fall time, contralateral, 30 dB SPL) was used to detect neuronal firing during the search phase. Upon encounter spiking activity of a single neuron, frequency response areas (FRA) and firing patterns to pure tones were characterized by presenting a range of different sound frequencies and intensities (100 ms duration, 5 ms rise/fall time, 0–100 dB SPL, 1-40 kHz, for both contra- and ipsi-lateral ear to the recorded IC); the different conditions were randomized in frequency and intensity within each protocol. Binaural response properties of neurons were tested by presenting pure tones at each neuron’s characteristic frequency (CF), while the contralateral ear being stimulated with the corresponding range of interaural intensity differences (±30 dB). Stimulus Specific Adaptation (SSA, Supp. Fig 3) was performed by first choosing 2 frequencies around the CF with 4 kHz difference (Duque and Malmierca, 2015a). Then both frequencies were exposed contralaterally in an alternating oddball paradigm, i.e. the first frequency was exposed 10% of the time while the other 90% and then the protocol was repeated exposing the first frequency more frequently while the second 10% of the time.

### Data acquisition

Recording electrodes were pulled from borosilicate glass capillaries (10–15 MOhm, BioMedical Instruments) with a PC-10 puller (Narishige) and then filled with 2 M NaCl solution, in which 2 % HRP (Type II, Sigma-Aldrich) was dissolved. First, the electrode was stereotactically aligned above the IC coordinates (Fig. 1), and then progressively lowered down in the tissue while presenting white noise to the animal. A careful approach to the recorded neuron guaranteed measurements of spikes from single neurons. In most parts of the IC, a crisp background response of Multi-Unit-Activity (MUA) to the noise stimulus could be monitored; recordings were considered to be single unit activity, when action potential amplitude was at least three fold the signal-to-noise ratio (snr) of the local MUA. At the end of each recording session the neighboring neurons were labeled with an injection of HRP by three 90 V DC current injections over 3 min (ISO-STIM 01D, NPI electronic).

**Figure 1:**
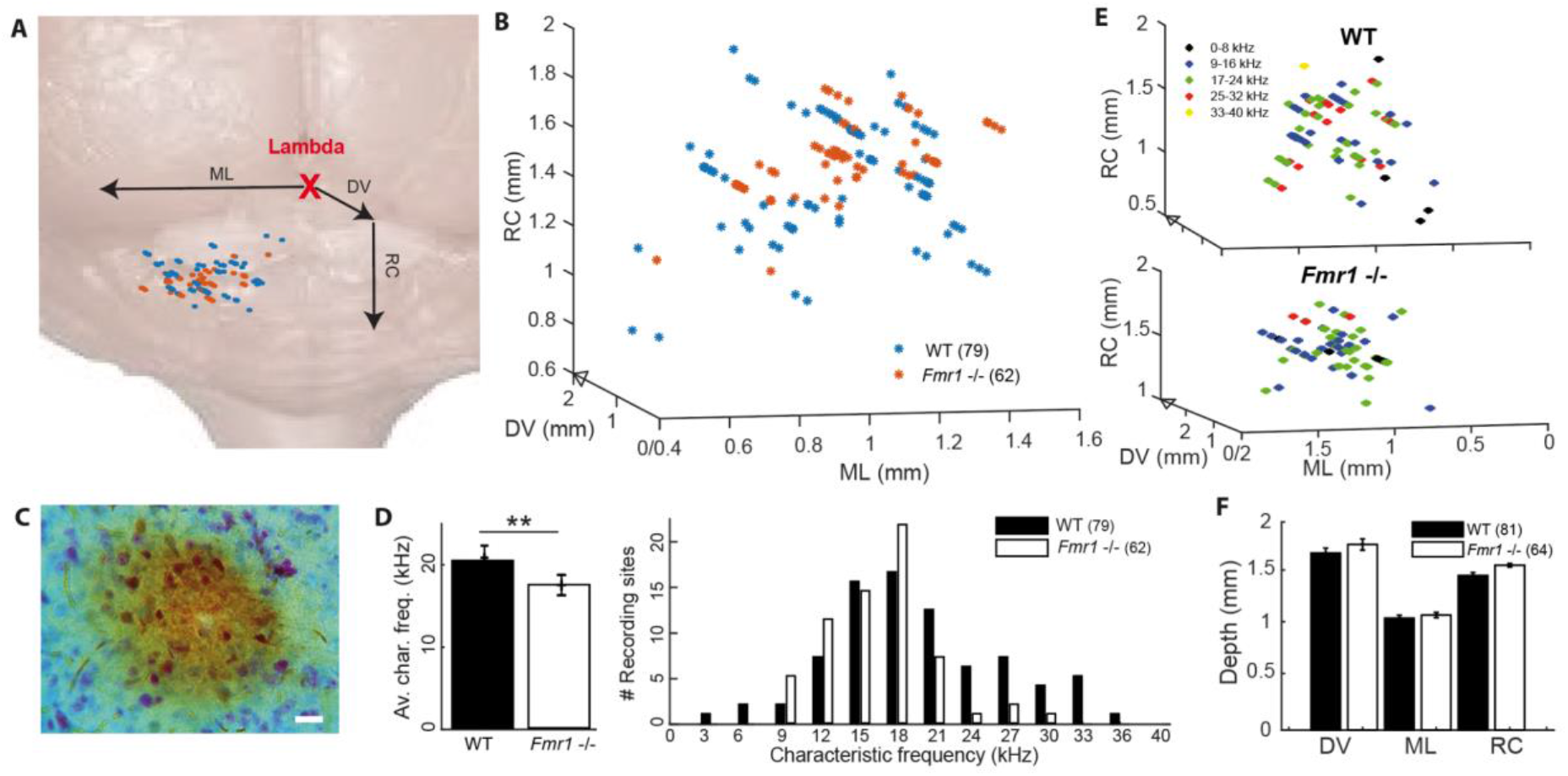
Location of the recorded IC neurons, with their characteristic frequencies. **A** Schematic reconstruction of stereotactic recording locations on top of a schematic brain picture, (orange) *Fmr1* -/- neurons, (blue) WT. **B** 3D-reconstruction of the different stereotactic locations. **C** Picture of an HRP staining (20µm). **D** (left) Histogram of all CF. (right) Averaged CF in the WT vs. *Fmr1* -/- mice. **E** 3D stereotactic locations represented with and color-coded CF (dark low, light color higher frequencies) in (up) all WT neurons, (down) all *Fmr1* -/- neurons.

### Stereotactic reconstructions

At the end of each experiment, animals were sacrificed with a lethal dose of ketamine/xylazine. The brain was removed and immersed in 4% PFA overnight. Brains were then thoroughly washed at room temperature with 0.1 M PBS, pH 7.4., and 70 µm coronal brain sections were cut on a vibratome (VT1200, Leica). HRP staining was done by incubating the sections in a solution containing Diaminobenzidine reaction (DAB kit, Sigma-Aldrich) and a Nissl counterstain. The pipette tracks, together with our stereotactic reconstructions (Sup. Fig 1), suggest that all neurons were recorded in the IC. All successful HRP staining confirmed that the recorded neurons were in the IC.

### Data analysis

Only single-unit recordings with a good signal to noise ratio (≥3) and stable firing over the entire time course of the experiment were kept for further analysis. For each neuron, a single compound PSTH was defined as the combination of the spiking activity averaged from all the different stimulus conditions screened during the FRA (0-100 dB, 1-40 kHz). This observation led us to establish a user-based distinction of four types of neurons: Transient neurons, Sustained neurons, Offset neurons and Inhibited neurons. This distinction was made using a simple two-steps decision tree looking at the FRA of each neuron and the compound PSTH: 1-Does the neuron fire mostly after the tone termination (in the offset period in its PSTH)? If so, this describes an Offset Neurons cf. Fig 2 D. In case of a negative answer then: 2-Does the FRA exhibit a negative tuning? This would be an "Inhibited neuron” cf. Fig. 2 C. Often, inhibited neurons have high spontaneous firing rates and their firing is reduced during tone exposure. If neither of these two conditions are matched -what is the case in about 80 % of the neurons of our dataset-then the neuron belong to one of the two more classical cell-types, the “Onset” or “Sustained” neurons, as broadly reported in the IC (Ehret et al., 2003; Gittelman et al., 2012). This user-based decision tree is the result of a very strong inhomogeneity of the physiology in our dataset and observed similarly in both genotypes.

**Figure 2:**
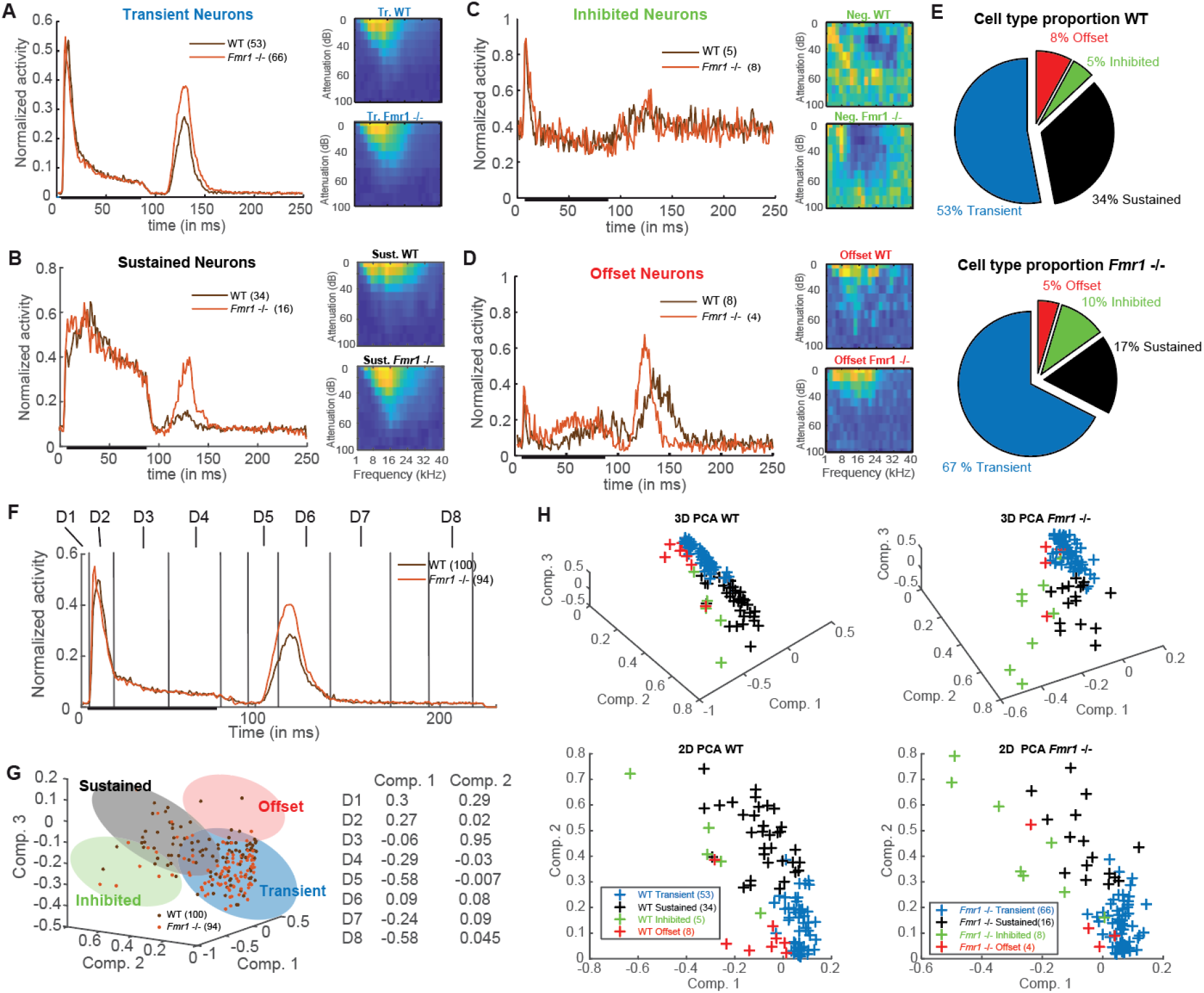
Four neuronal response patterns in both genotypes. **A** (left) Averaged compound PSTH of all transient neurons, (brown) WT and (orange) *Fmr1* -/-; (right above) Averaged FRA of all WT transient neurons and (right below) FRA of all *Fmr1* -/- transient neurons. **B** (left) Averaged PSTH of all sustained neurons WT, and *Fmr1* -/-; (right above) FRA of all WT sustained neurons, and (right below) their *Fmr1* -/- counterparts. **C** (left) PSTH of all inhibited neurons, WT and *Fmr1* -/-; (right above) FRA of all WT inhibited neurons, and (right below) their *Fmr1* -/- counterparts; note the negative tuning that these neuron’s FRA exhibit. **D** (left) PSTH of all offset neurons, WT and *Fmr1* -/-; (right above) FRA of all WT all offset neurons, and (right below) their *Fmr1* -/- counterparts. **E** Proportion of each cell types in (above) WT and (below) *Fmr1* -/- neurons. **F** All-cell average of all compound PSTHs in the WT and *Fmr1* -/- mice. Different values of the transient firing are taken in each time-window for each cell (D1-D8), used for the following principal component analysis (PCA). **G** (left) 3D-representation of all recorded neurons along the first three components. Note the four shaded areas corresponding to the four cell types detailed above. (right) Projections of the two first principal components on the eight time-windows. **H** (above left) 3D-representation of all WT neurons; (below left) same neurons represented along the two first components; (above right) 3D-representation of all the *Fmr1* -/- neurons; (below right) 2-D projection of all the *Fmr1* -/- neurons on the two first component.

To address the validity of such a distinction we performed a few controls, i.e. a Principle Component Analysis (PCA) to illustrate the diversity of the compound PSTHs, a comparison of stereotactic depth and a more detailed PSTH comparison for FRA sub-sections. The PCA was based on the average spiking rate in eight different time windows (Dimension 1: 0-5 ms, Dim. 2: 5-20 ms, Dim. 3: 20-50 ms, Dim. 4: 50-90 ms, Dim. 5: 95-110 ms, Dim. 6: 110-140 ms, Dim. 7: 140-170 ms, Dim. 8: 190-230 ms).

### Quantification of frequency response area (FRA)

The CF was defined as the frequency at which each neuron responded (firing 3x-larger than spontaneous activity) to the lowest intensity (threshold). Sharpness of tuning was analyzed as previously published (Garcia-Pino, et al. 2017) and reported in Q10, 30, 50 values (Fig. 3); Q10 values were calculated as the response bandwidth at 10 dB above each neuron’s threshold divided by the CF; Q30 and Q50 were calculated accordingly at 30 dB and 50 dB above each neuron’s threshold.

**Figure 3:**
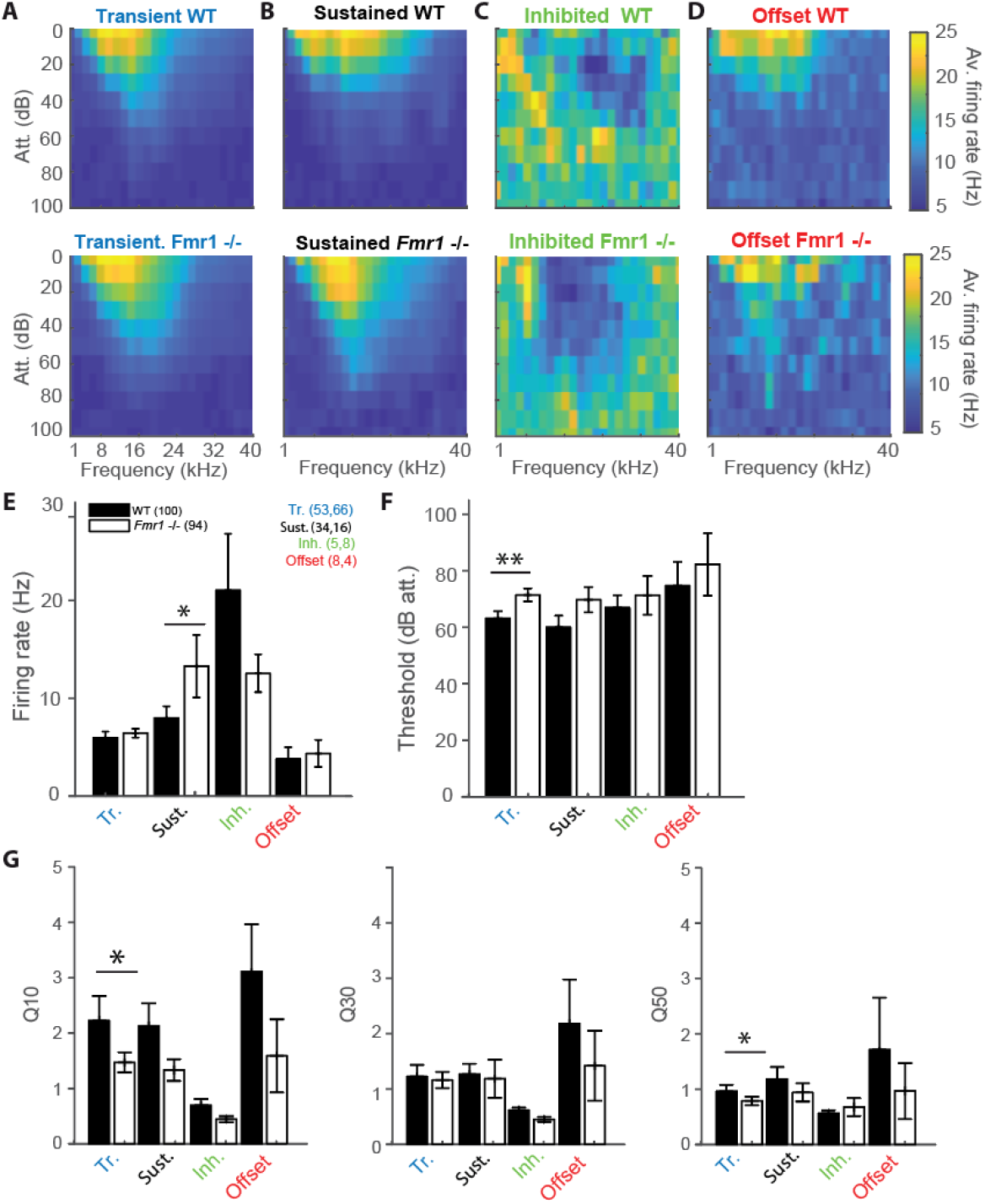
*Fmr1* -/- neurons are over-activated and oversensitive. **A** (above) Averaged of all WT transient neurons FRA, and (below) all their *Fmr1* -/- counterparts. **B** (above) Averaged of all WT sustained neurons FRA, and (below) their *Fmr1* -/- counterparts. **C** (above) Averaged of all WT inhibited neurons FRA, and (below) their *Fmr1* -/- counterparts. **D** (above) Averaged of all WT offset neurons FRA, and (below) their *Fmr1* -/- counterparts. **E** Averaged firing rate of each of the 4 classes. **F** Average threshold of the 4 classes together. **G** Averaged Q values measured in each neuron, resp. (left) Q10, (middle) Q30, (right) Q50.

### Further data analysis

First spike latency was measured at the CF (Fig. 4) as the time point following tone exposure when the average spiking of each neuron reached three times its own spontaneous firing baseline, measured in the 5 ms before tone exposure. Inhibited and offset neurons were not reported as their spiking patterns do not exhibit such measurable increases: Offset neurons barely have tone modulations at the beginning of the tone, inhibited neurons have a decreasing firing concomitant to the activation of the other neurons, suggesting a possible inhibitory modulatory role. The time of maximum firing was measured at the peak of the compound PSTH. Rebound activity was characterized by averaging the firing rate over a time window following tone exposure (100 to 150 ms). The Interaural-Level-Difference (ILD) tuning curve was obtained by gathering the different ipsilateral and contralateral stimulus intensities according to their respective ILD values. Once a neuron’s CF and threshold was identified, a set of different intensities were presented (i.e. threshold/-10/-20/-30 dB) with contralateral intensities ranging between -30 to +30 dB ILD for each ipsilateral intensity. The spikes elicited by each neuron were summed up and normalized to the different ipsilateral intensities within a fixed ILD window (−30 to +30 dB) to produce the estimated spike rate per cells for each different ILD. Once averaged the obtained ILD curve was then characterized by the level of the half-maximum and by the overall slope steepness measured from the maximum to the minimum of the obtained normalized ILD curve. Finally, the adaptive behavior of neurons to novel stimuli was measured by performing mismatch of two pure tones (Sup. Fig. 3 A), choosing two frequencies ± 4 kHz around the CF at 30 dB below the threshold in a 10-90 % alternation paradigm (Ulanovsky et al., 2003; Malmierca et al., 2009).

**Figure 4:**
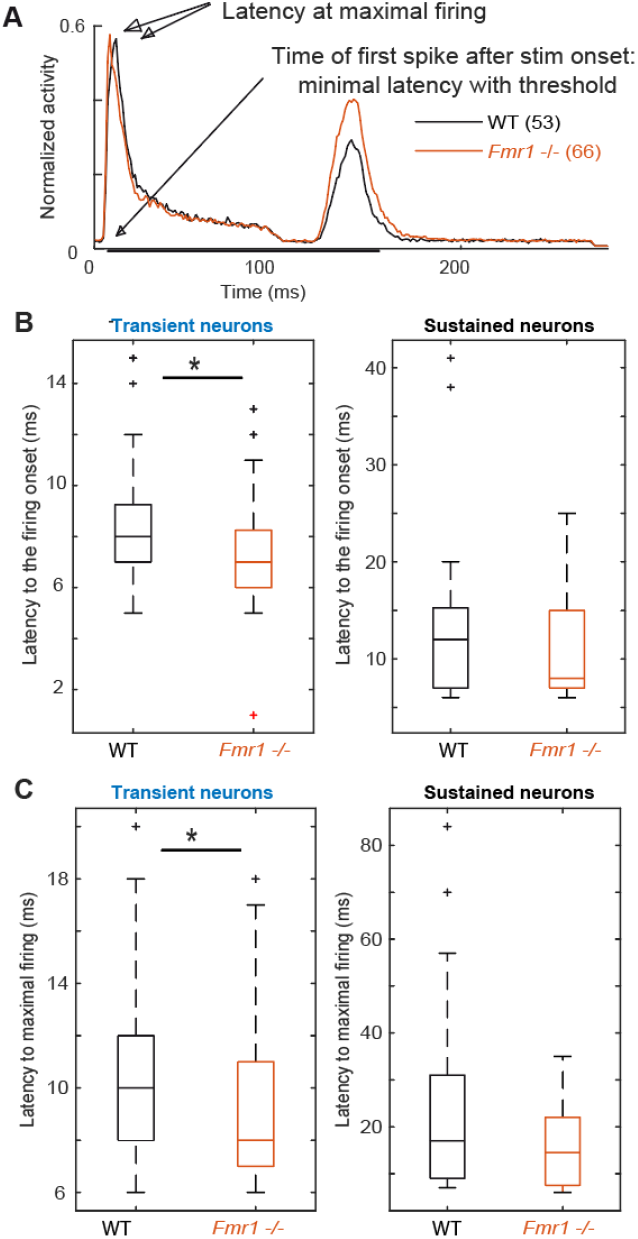
*Fmr1* -/- neurons exhibit an earlier and stronger activation. **A** Averaged and normalized PSTH of all cells over all conditions during the FRA for all standard WT and *Fmr1* -/- neurons. **B** Boxplot representation of the averaged latency in transient and sustained neurons. **C** Boxplot representation of the timing of maximal spiking in transient and sustained neurons from WT and *Fmr1* -/- mice.

## Results

### Higher frequencies are under-represented in the Inferior Colliculus of the Fragile X mouse model

A recent study provided evidence that sound representation is altered in IC neurons of developing *Fmr1* -/- mice (Nguyen et al. 2020), however, it is unclear whether these differences are maintained in fully mature mice. To do so we presented 100 ms long tones of different frequencies (1-40 kHz, 25 steps) and different sound intensities (0-100 dB att., 10 dB steps) to the contralateral ear. From this FRA the CF of each neuron, the frequency where a neuron responds to the lowest intensity, was extracted (Fig 3 A-D). The CF was correlated with the location of each neuron within the IC as confirmed by HRP staining (Fig.1A, see methods) and subsequent reconstruction according to the stereotactic coordinates. Neurons were located in the entire inferior colliculus, including the dorsal and external cortices of the IC (Fig. 1F). The average location of the recorded neurons did not differ between genotypes in either of the three axes of the micromanipulator arm in our reference frame (Fig. 1 B-E-F, Sup. Fig. 1). Nevertheless, neurons of WT mice responded best to significantly higher frequencies compared to neurons in *Fmr1* -/- mice (Fig. 1 D left, 20.43 kHz ± 0.77 vs 17.48 kHz ± 0.56, unpaired t-test, P = 0.0074). In fact, CF higher than 24 kHz were almost absent in *Fmr1* -/- mouse IC (Fig 1 D right) despite being recorded in similar if not deeper depth (Sup. Fig. 1). Whether a loss of FMRP impacts tonotopic representation in the IC, as previously shown in the MNTB (Strumbos et al. 2012, Mccullagh et al. 2017), needs to be confirmed using different methods.

### Four different response patterns of IC neurons in both genotypes

Previous studies reported broader tuning, higher thresholds, increased spiking activity and latency changes in the ascending auditory pathway of adult *Fmr1* -/- mice (Brown et al. 2010, Stumbos et al. 2019, Garcia-Pino et al. 2017). Analyzing these parameters across our dataset of all recorded IC neurons did not yield any statistical differences between WT and *Fmr1* -/- mice. Using molecular, anatomical and physiological markers the auditory midbrain is composed of different neuron types, however a comprehensive neuron type classification combining various methodologies has not been achieved in this brain region (Wallace et al. 2021). Moreover, extracellular spike shape analysis, which is widely used in other brain regions for neuron type classification, has not been proven successful in the IC (Ito, 2020; Wallace et al., 2021). We therefore classified IC neurons, as previously done (Tan et al., 2007; Palmer et al., 2013; Akimov et al., 2017), according to their spike response patterns to pure tone stimulation with different frequencies using a simplified decision tree (see methods). As response patterns change depending on the frequency and intensity of the sound, we used a compound PSTH defined as the combination of all the spiking patterns gathered from all different conditions. The averages of these compound PSTHs for each category of the different neuron types are shown in Fig. 2A-C. Transient neurons, which are present in both genotypes, are characterized by transient responses to the onset of the tone (Fig. 2 A). In many cases they show an offset-rebound activity following the termination of tone exposure (Fig. 2 A left). Both genotypes exhibit classical frequency tuning (Fig. 2 A right). Those neurons represent the majority of the recorded neurons (67 % in the *Fmr1* -/-, 53 % in the WT, Fig. 2 E out of resp. 94 *Fmr1* -/- neurons and 100 WT neurons) and likely cover the previously reported “ON”, “Long-Latency” and “ON-OFF” neurons from other studies (Duque et al., 2012; Palmer et al., 2013; Akimov et al., 2017). They may also overlap with the “Sensitive-symmetrical”, the “Sensitive-asymmetrical” and the “Selective” neurons from other authors (Lee et al. 2019). Second in proportion, the sustained neurons exhibit a more regular firing during the entire duration of sound exposure (Fig. 2 B left). They also exhibit a more classical excitatory frequency tuning (Fig. 2 B right). The sustained neurons are more abundant in the WT compared to *Fmr1* -/- mice (34 % vs 17 % Fig. 2 E). This neuronal type likely covers the “ON-Sustained” and “Sustained” neuron type from other studies (Duque et al., 2012; Palmer et al., 2013; Akimov et al., 2017). Third, inhibited neurons exhibit the very peculiar “negative tuning” characteristic of their FRA (Fig. 2 C right); they have a negative firing modulation during tone exposure (Fig. 2 C left). This neuron type exhibits a high spontaneous firing compared to the other types; it represents a minority of 10 % of *Fmr1* -/- and 5 % of WT neurons. This neuron type matches very well the already reported “inhibitory frequency response area” with un-classical FRA; they were initially reported to be exclusively present in awake mice (Duque and Malmierca, 2015a) as well as the inhibited neurons in the IC’s cortex (Wong and Borst, 2019). Fourth, offset neurons exhibit the largest response after the termination of the sound exposure (Fig. 2 D left); they exhibit a more usual FRA (Fig. 2 D right). These neurons represent 8 and 5 % of the whole dataset in resp. WT and *Fmr1* -/- neurons. All further analysis of response characteristics were performed using these groups.

To address the validity of such a distinction based on a simple decision tree, we controlled for few aspects supporting this categorization: first, we projected the PSTH of each cell, showing their discrepancy of firing pattern using Principal Component Analysis (PCA) in both genotypes (Fig. 2 F-G-H). Our PCA was based on average spiking rate in eight different time windows (Fig. 2 F). The values decomposition of the first principal component having the highest weight (Fig 2G) are based on the transient onset (Dim. 2), the sustained firing (Dim. 3-4) and the offset responses (Dim. 5, Dim. 7-8). An unexpected outcome of this PCA is the similar clustering of the four neuronal cell types observed in both WT and *Fmr1 -/-* mice (Fig. 2 G- H). Interestingly, the WT transient neurons (blue) seem to be composed of two distinct neuronal populations, which we were unable to further dissect in a meaningful way (Fig 2 H bottom). Second, we controlled that the stereotactic locations of the recorded neurons were not accounting for neuron type in either depth or location (Sup. Fig. 1 A-B). These data suggest that the different cell types were pooled from a homogenous region and were not the result of differences in the physiology of IC cortex versus IC central nucleus neurons. Third, to prevent any bias resulting from oversampling of PSTHs for multiple tone frequencies or intensities, we split the conditions recorded during the FRA into different features (all intensity at one frequency, all frequency for one intensity, etc.) and observed consistent differences in spiking patterns according to the distinguished cell-types in both genotypes (Sup. Fig. 2). Interestingly, we consistently observed offset responses in all cell types when gathering spiking outside their excitatory tuning area (Sup. Fig. 2).

### *Fmr1* -/- neurons have lower thresholds, broader tuning and shorter response latencies compared to WT neurons

We reexamined thresholds, firing rates, frequency tuning and response latencies of different IC neuron types in WT and *Fmr1* -/- mice. The class-averages of all FRAs within each neuron type exhibited similar tuning between the different genotypes (Fig. 3 A-B-C-D). Nevertheless, when processing each neuron type individually we revealed a clear bias of lower thresholds in all neuron types of *Fmr1* -/- mice (Fig. 3 E-G). This bias reached significance levels only in transient neurons (Fig 3 E, 71.66 dB ± 2.3 in the *Fmr1* -/-, 63.39 dB ± 2.54 in the WT, unpaired t-test P=0.0173). All other neuron types exhibited lower thresholds in *Fmr1* -/- mice but failed to pass significance probably due to the limited sampling. Sustained *Fmr1* -/- neurons showed significantly higher firing rates than their WT counterparts (Fig. 3 F, 13.27 Hz ± 2.6 in *Fmr1* -/-, 7.97 Hz ± 1.19 in WT neurons, P=0.0255). The firing rates in other *Fmr1* -/- neuron types were also higher (Fig. 3 F), except for the *Fmr1* -/- inhibited neurons. Taken together, these results indicate a generally increased firing in the *Fmr1* -/- mice, which matches the broadly reported hyper-active, hyper-sensitive phenotype of neuronal sound representation in FXS mice.

We next analyzed frequency tuning width of each neuron using the Q10, Q30 and Q50 values as a measurement for tuning widths, with lower Q values indicating broader tuning (see methods). Surprisingly, only transient neurons showed significantly broader tuning for Q10 values (Fig 3 G, in the transient neurons 1.45 ± 0.18, in the *Fmr1* -/-, 2.23 ± 0.44 in the WT, P=0.0495) and Q50 values (0.74 ± 0.07, in the *Fmr1* -/-, 0.80 ± 0.10 in the WT, P=0.0437). In all other neuron types tuning width was similar for both genotypes.

Analyzing first spike latency and the latency of maximal firing revealed shorter latencies for transient and sustained neurons of *Fmr1* -/- mice (Fig. 4). Transient and sustained *Fmr1* -/- neurons exhibit a significant shorter first spike latency (Fig. 4. B, 7.60 ms ± 0.27 in the *Fmr1* -/- vs 8.53 ms ± 0.35 in the WT for transient cell, unpaired t-test P = 0.0489; 11.36 ms ± 1.85 in the *Fmr1* -/- vs 13.81 ms ± 1.64 in the WT for sustained cell, unpaired t-test P = 0.0709). They also exhibit significantly earlier maximal firing (Fig. 4. C, 9.42 ms ± 0.36 in the *Fmr1* -/- vs 10.36 ms ± 0.44 in the WT for transient cell, unpaired t-test P = 0.0426; 15.7 ms ± 2.22 in the *Fmr1* -/- vs 23.05 ms ± 0.39 in the WT for sustained cell, unpaired t-test P = 0.3310).

Taken together, these results suggest that *Fmr1* deletion has similar, but not larger effects on spike number and tuning width compared to auditory brainstem neurons.

### Enhanced offset-rebound activity in sustained and transient neurons of *Fmr1* -/- mice

Unexpectedly, post-stimulus or offset-rebound activity differed between *Fmr1* -/- mice and WT neurons (Fig. 5). In both genotypes and in both transient and sustained neurons offset- rebound activity was present (Fig. 5 A -B). However, more neurons exhibited offset-rebound activity in *Fmr1* -/- compared to WT (Fig. 5 A -B). Quantification of offset-rebound firing yielded significant higher values for *Fmr1* -/- compared to WT mice (Fig 5 C, left: 0.53 Hz ± 0.03 in *Fmr1* -/- 0.37 Hz ± 0.04 in the WT transient neurons, unpaired t-test P = 0.0014, 0.46 Hz ± 0.07 in *Fmr1* -/- 0.26 Hz ± 0.04 in the WT sustained neurons, unpaired t-test P = 0.0254, and the corresponding histogram Fig. 5 C, right). It is also noticeable that in the *Fmr1* -/- mice’s IC, almost all transient neurons exhibited rebound activity (60/66), while in the WT only about two third of the neurons show an offset reponse (39/54). Furthermore, the tuning exhibited by the rebound activity was always “negative” (Fig. 5 D) and is most likely related to “inhibitory frequency response areas” as characterized by other authors (Duque and Malmierca, 2015b). Our results indicate that most IC neurons (transient & sustained) exhibited a classical FRA firing to pure tone exposure, which is characterized by spiking activity during the stimulus presentation. In many transient neurons, this is then followed by an offset-rebound firing whose tuning is outside of the excitatory tuning area (Fig 5 E). Similar differences were also observed in sustained neurons (Fig 5 E), arguing for a cell-type independent mechanism. An increased offset-rebound activity could potentially arise from a change in each neuron’s intrinsic ion channel composition and/or an enhanced lateral inhibition (Kopp-Scheinpflug et al., 2018). This indicates that not only excitation but also inhibition is homeostatically upregulated in *Fmr1* -/- mice.

**Figure 5:**
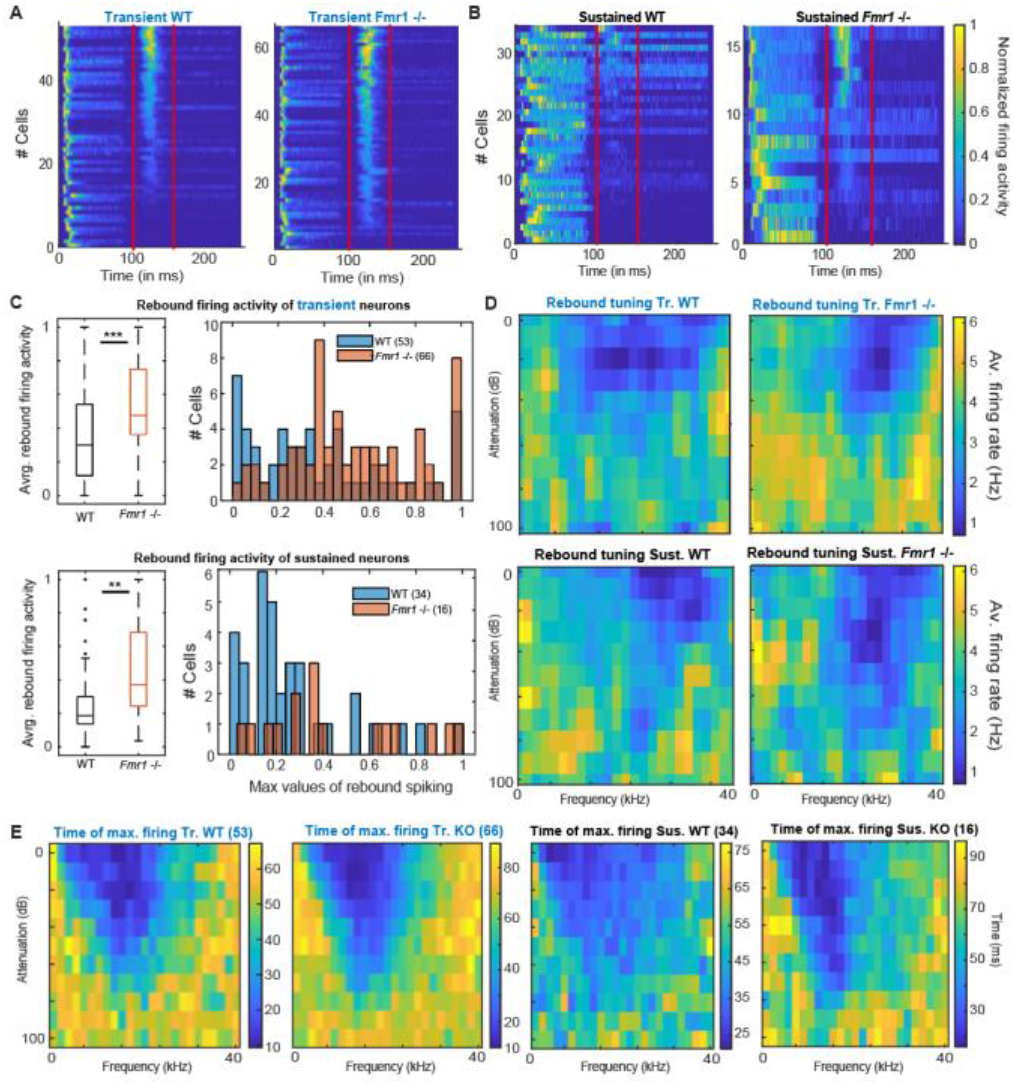
Offset-rebound activity in *Fmr1* - /- neurons. **A** Normalized PSTH for (left) all WT standard neurons and (right) their *Fmr1* -/- counterparts; note that the red lines indicate the limits used to quantify the tuning of the rebound-offset activity. **B** Normalized PSTH (left) all WT sustained neurons and (right) all their *Fmr1* -/- counterparts. **C** (above left) Averaged and normalized rebound-offset activity in all WT and *Fmr1* -/- standard cells; (above right) Corresponding histograms, note the semi- Gaussian organization of the *Fmr1* -/- offset- rebound; (below left) Averaged and normalized rebound-offset activity in all WT and *Fmr1* -/- sustained neurons; (below right) Corresponding histogram. **D** Averaged normalized FRA from spikes selected from offset-rebound activity which illustrates the tuning exhibited by the rebound-offset activity in respectively: (above left) all WT standard neurons, (above right) all *Fmr1* -/- standard neurons, (below left) all WT sustained neurons, (below right) all *Fmr1* -/- sustained neurons. **E** Similarly for each single individual frequency and attenuation, the time of maximal firing is averaged within the populations and illustrated in the corresponding color code to report the later peak in the PSTH outside of the FRA for (left) Transient WT, (middle left) transient *Fmr1* -/-, (middle right) sustained WT and (right) *Fmr1* -/- neurons.

### Binaural balance of excitation and inhibition (EI-balance) is shifted in *Fmr1* -/- mice

Our previous results using only monaural sound stimulation indicate both an enhancement of excitation within the excitatory tuning area and possibly, an increased inhibition resulting in rebound spiking outside the excitatory tuning area. Using binaural stimulation our previous study in the LSO revealed enhanced ipsilateral, excitatory and no change in contralateral, inhibitory input strength in *Fmr1* -/- mice thereby shifting the responses to binaural stimulation towards the ipsilateral ear (Garcia-Pino et al., 2017). We were interested in whether these binaural response properties are preserved at the level of the IC or whether additional homeostatic mechanisms may compensate or enhance the changes observed in *Fmr1* -/- LSO neurons. Accordingly, we presented a set of different sound intensities at the ipsi- and contralateral ear. Spike responses with sounds of different intensities but same ILDs were added and normalized to the maximal responses. Fig. 6A displays the response levels to these added ILDs for each cell. Only neurons with a sustained and transient response were included in the analysis (83 out of 100 WT neurons and 84 out of 94 *Fmr1* -/- neurons), but neurons were not selected for binaural sensitivity. Interestingly the summed binaural responses of all WT neurons revealed a strong bias with excitation predominating for contralateral stimulation and inhibition for increasing ipsilateral stimulation. This was not the case in *Fmr1* -/- neurons, where ILD functions were much flatter and ipsilateral stimulation suppressed contralateral evoked spiking much less (Fig. 6A-C). This was observed in both sustained and transient firing neurons.

**Figure 6:**
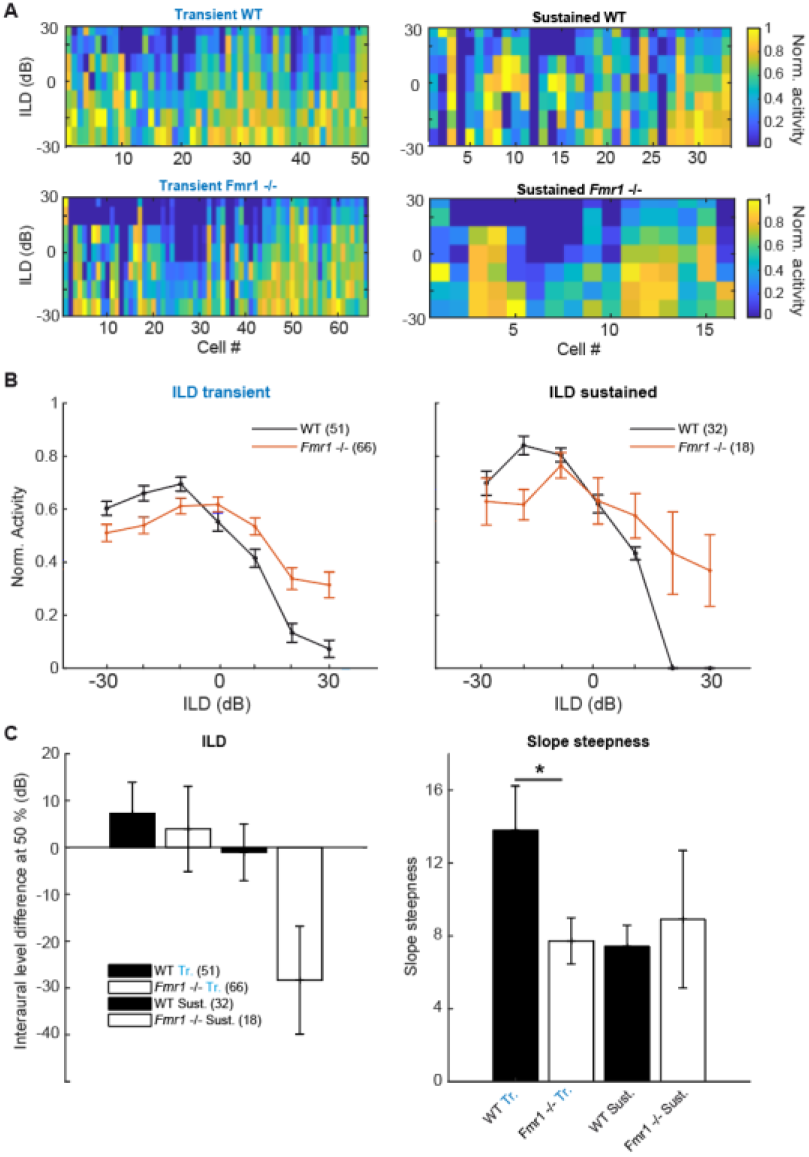
Binaural integration of ILDs is impaired in the *Fmr1* -/- mice. **A** Normalized spike rate per cell for the different ILD conditions for: (above left) all WT standard neurons; (below left) all *Fmr1* -/- standard neurons; (above right) all WT sustained neurons and (below right) all *Fmr1* -/- sustained neurons. **B** (left) Averaged normalized ILD for all standard neurons WT and *Fmr1* -/-; (right) ILD for all sustained neurons WT and *Fmr1* -/-. **C** Quantification of resp. (left) averaged ILD as the half height of the corresponding fitted sigmoid; (right) for the slope steepness of resp. WT and *Fmr1* -/- ILDs.

In transient neurons we observed a small shift of the ILD function in *Fmr1* -/- mice in combination with a stronger firing for ILDs larger than 20 dB. In both transient and sustained neurons, a clear difference of ILD turning point was observed (Fig. 6C) and a significant difference in the slope was seen in the transient neurons (*Fmr1* -/- 7.72 ± 1.27 vs WT 13.79 ± 2.45, unpaired t-test P = 0.0457). In the sustained neurons we observed a similar difference also indicating a shift in the balance of excitation and inhibition. This suggests that the imbalance of binaural excitation and inhibition, as observed in the LSO, is mostly preserved at the level of the IC. Whether this also holds for stimuli with more complex sound features remains to be shown.

### Weak stimulus specific adaptation in the IC of WT and *Fmr1* -/- mice

People with Fragile X syndrome show decreased processing abilities of novel stimuli and less habituation when presented with an auditory odd-ball paradigm (Ethridge et al., 2020). Neurons in the IC are known to exhibit an enhanced representation to pure tones when they are exposed in an oddball fashion. This is usually measured by choosing two tones around a neuron CF (Duque and Malmierca, 2015). In addition, the upper or the lower frequency is exposed irregularly (10% randomized) inducing an increased firing compared to the frequent exposure. In our hands none of the two probed frequency Specific adaptation Indices (SI) reached the expected averaged positive values (Sup. Fig. 3 D). A few neurons exhibited meaningful adaptation, but no group of neurons showed a reliable oddball adaptation. The Common Stimulus specific adaptations index (CSI) using responses to both frequencies showed a similar profile (Sup. Fig. 3 D). Overall, our results tend to confirm the existence of adaptation in the IC (Sup. Fig. 3 C), but do not report reliable effects at the population level.

## Discussion

We studied classical features of IC neuron responses presented with simple sounds (pure tones) to obtain further insights into the pathophysiological basis of distorted auditory processing in FXS. One major finding that has not been previously described was that the post-inhibitory rebound response, mainly evoked by stimulating outside of the excitatory receptive field, was substantially enhanced in neurons from *Fmr1* -/- mice. All other response features such as total spike number, tuning width and binaural integration were similar to the ones observed in lower brainstem nuclei. Due to the diversity of IC neuron types, we observed statistical differences between WT and *Fmr1* -/- mice, only once IC neurons had been categorized according to their response patterns to pure tones of different frequencies and intensities.

### Categorization of IC response types is similar in WT and *Fmr1* -/- mice

Comparing neuronal responses to pure tones between WT and *Fmr1* -/- mice in all neurons did not yield substantial differences. As a consequence, we grouped the neurons based on their firing patterns to short pure tones of different frequencies and intensities, and on whether neurons were excited or inhibited by sound stimulation. Similar classes of response patterns in the IC have been suggested in various species including bats and mice (Xie et al., 2007; Palmer et al., 2013; Akimov et al., 2017). The transient and sustained neurons compose most of our dataset, which matches most of the published literature, whereas offset neurons and inhibited neurons remain a minority in our recordings and in the published literature (Willott et al., 1988; Kasai et al., 2012; Wong and Borst, 2019). One reason for the low number of inhibited neurons reported so far could be that measurement of inhibited neurons is tricky. As these neurons exhibit a very high spontaneous firing rate, they can be miss-identified as dying neurons.

In previous studies classification of firing types in IC neurons was mostly based on the firing pattern to a single frequency and intensity, mostly at the CF (Nasimi and Rees, 2010; Gourévitch et al., 2020) or, as in other studies, by the shape of the excitatory and/or inhibitory frequency response areas (Palmer et al., 2013). With our approach we included information on active inputs across the entire FRA into the classification scheme, by using the summed spiking activity to all presented frequencies and intensities similarly to a previous study (Egorova et al., 2020). This includes not only information on the excitatory receptive fields, but also on inhibitory receptive fields. This inhibition has been shown to induce rebound activity after sound termination in many cases (Kopp-Scheinpflug et al., 2018). We then used a simple user-based decision tree to categorize spiking patterns, which was followed by PCA confirmation. For larger datasets, such a PCA representation could be used to cluster the neuronal response types to replace the user-based decision tree. However, in our case it would lack reproducibility due to the relatively small sample size. Very recently, several new methods have been developed to categorize neuron types based on spike patterns. This includes dimensionality reduction algorithms such as UMAP (Lee et al., 2021) or t-SNE (Dimitriadis et al., 2018). However, these algorithms require larger datasets, as for example acquired with high-density silicon probe recordings.

Our data also indicate that *Fmr1* deletion decreases cell type diversity as transient neurons represent 53% of all neurons in the WT and 67% in the *Fmr1* -/-. Furthermore, the representation of response features of each neuron in both WT and *Fmr1* -/- using PCA illustrates that *Fmr1* -/- neurons exhibit less clear boundaries between the different neuron types compared to WT neurons, where transient neurons exhibit a more compact cloud of points. Up to now correlating spiking response patterns or spike shape of extracellularly recorded IC neurons with other molecular, cellular or morphological features has not been successful (Ito, 2020; Wallace et al., 2021). More studies on this matter are needed to obtain a better understanding of IC network functions.

### Genotype specific changes in firing rate, frequency tuning and binaural response properties in IC neurons are similar to auditory brainstem neurons

We observed a significant increase in firing rate in *Fmr1* -/- mice only in sustained neurons. As the proportion of transient to sustained firing neurons also increased in *Fmr1-/-*, the interpretation of this result is difficult. It is possible that firing patterns have changed upon gene deletion resulting in different categorization of neurons. Nevertheless, these data indicate that a transition of onset or transient firing neurons in WT to sustained firing neurons in *Fmr1* -/- mice, as previously shown in the LSO (Garcia-Pino et al., 2017), is not occurring in the IC. Similar increases in IC firing rate have been reported in previous studies and were either small (Mott and Wei, 2014) or selective to neurons with CF below 20 kHz (Nguyen et al., 2020). Taken together, the data suggest that firing rates in *Fmr1* -/- mice are not substantially increasing between the auditory brainstem and IC, possibly due to homeostatic compensatory mechanisms being activated. Recordings from awake behaving mice may shed further light on the correlation of neuronal activity patterns with the clear acoustic hypersensitivity observed in FXS (Sinclair et al., 2016).

Similar to previous studies we measured broader frequency tuning in *Fmr1* -/- neurons (Mott and Wei, 2014; Nguyen et al., 2020), but tuning width, as quantified by Q-values, was only significant in transient firing neurons. Tuning bandwidth in *Fmr1* -/- LSO neurons is also significantly wider, which we suggested to be a reflection of reduced pruning of excitatory inputs during development (Garcia-Pino et al., 2017). Since frequency response tuning in the IC is not a mere projection from the auditory brainstem but reshaped by the interaction of excitatory and inhibitory inputs (Alkhatib et al., 2006; Lee et al., 2019), the broadening of tuning in *Fmr1* -/- neurons may also occur in the IC directly.

Our data also suggest that ILD tuning in the IC is severely affected by *Fmr1* deletion with ipsilateral evoked inhibition being weakened. This is different to the changes in ILD representation in the LSO of *Fmr1* -/- mice where ipsilateral excitation was exaggerated whereas contralateral inhibition was largely unaffected (Garcia-Pino et al., 2017). One possible explanation may be that in adult animals, inhibition to LSO neurons is transmitted by glycine, whereas in the IC GABA receptors predominate. Indeed, GABAergic, parvalbumin expressing neurons or reduced GABA transmission was observed in cortical circuits of *Fmr1* -/- mice (Selby et al., 2007; Song et al., 2022). Many binaural features in the IC are either refined or even created de novo in the IC (Park et al., 2004; Pollak, 2012), and GABAergic inhibitory inputs either from the dorsal nucleus of the lateral lemniscus or from the opposite IC may be impaired by *Fmr1* deletion.

In general, response features of IC neurons in *Fmr1* -/- are overall not much different to neurons in the auditory brainstem indicating homeostatic compensatory mechanisms to be effective and thus preventing the development of exaggerated spike responses in the IC. These compensatory mechanisms seem to be activated later during development, since response increases in the IC are largest in juvenile *Fmr1* -/- animals and decreases again once animals have reached adulthood (Nguyen et al., 2020).

### Enhanced activity to the offset of a sound in *Fmr1* -/- mice

An interesting feature we found was an increased offset response in *Fmr1* -/- mice in both transient and sustained firing neurons. Generally, largest offset responses were elicited in both genotypes, when neurons were stimulated above or below their excitatory frequency response area, where inhibitory inputs predominate. Indeed, offset spiking, defined as spiking activity in response to the end of a sound stimulus, has been reported in the IC (Tan et al., 2007; Kasai et al., 2012) and in other nuclei of the auditory brainstem, especially in neurons of the superior paraolivary nucleus (Felix et al., 2011; Kopp-Scheinpflug et al., 2011). In the IC, offset activity may be evoked by excitatory synaptic inputs occurring after sound termination or by inhibitory rebound activity (Kasai et al., 2012). The temporal pattern of the excitatory and inhibitory responses to the beginning and the end of a sound stimulus also determines the selectivity of neurons to the duration of a sound (Ono and Oliver, 2014; Valdizón- Rodríguez and Faure, 2017; Valdizón-Rodríguez et al., 2019). What are the possible underlying cellular mechanisms associated with this enhanced offset activity in *Fmr1* -/- neurons? Rebound spiking from inhibition depends on the interplay of inhibitory inputs in combination with the activation of Hyperpolarization-activated and Cyclic Nucleotide gated (HCN) channel and in many cases low-threshold calcium-channels (Kopp-Scheinpflug et al., 2011, 2018). Enhancement in any of these ionic currents may lead to the increased offset activity in *Fmr1* - /- mice. However, previous studies on inhibitory synaptic transmission indicate a reduced inhibitory transmission and not an enhanced inhibitory transmission (Cellot and Cherubini, 2014), whereas HCN channel activity can be modulated in both directions by *Fmr1* deletion (Brager et al., 2012; Brandalise et al., 2020). It is possible that this altered presentation of sound termination in auditory neurons of *Fmr1-/-* contributes to the observed deficits in the processing of speech and communication sounds in humans with FXS and in Fragile X animal models (Engineer et al., 2014; McCullagh et al., 2020).

## Conclusions

Despite almost three decades of research on the Fragile X mouse model, we still fail to precisely understand where and how the genetic malformation does impair the brain, and its auditory system in particular. Recent evidence suggests that the IC plays a predominant role in FXS as the deletion of *Fmr1* -/- expression specifically in VGlut2 neurons both the cortex and the IC is sufficient to induce audiogenic seizure, while IC Ntsr1 specific *Fmr1* deletion is necessary but not sufficient to induce audiogenic seizure in the Cortex (Gonzalez et al., 2019).

Regarding this fact, it is surprising to see spike rates in IC neurons of *Fmr1* -/- mice being only moderately elevated, considering how strong the acoustic behavior of FXS patients and mice is affected. The larger and faster excitation seems to be present at every stage of the ascending auditory system, however, upregulation of inhibition, probably by homeostatic regulation, may come into place at the level of the IC and prevent IC neurons from overexcitation. These compensatory mechanisms may, however, directly influence spike patterns, as we observed with the enhanced offset activity to the end of a sound stimulus. To what extent this impairs the analysis of complex sound needs to be addressed in future studies.

## Supplementary figures

**Supplementary figure 1:**
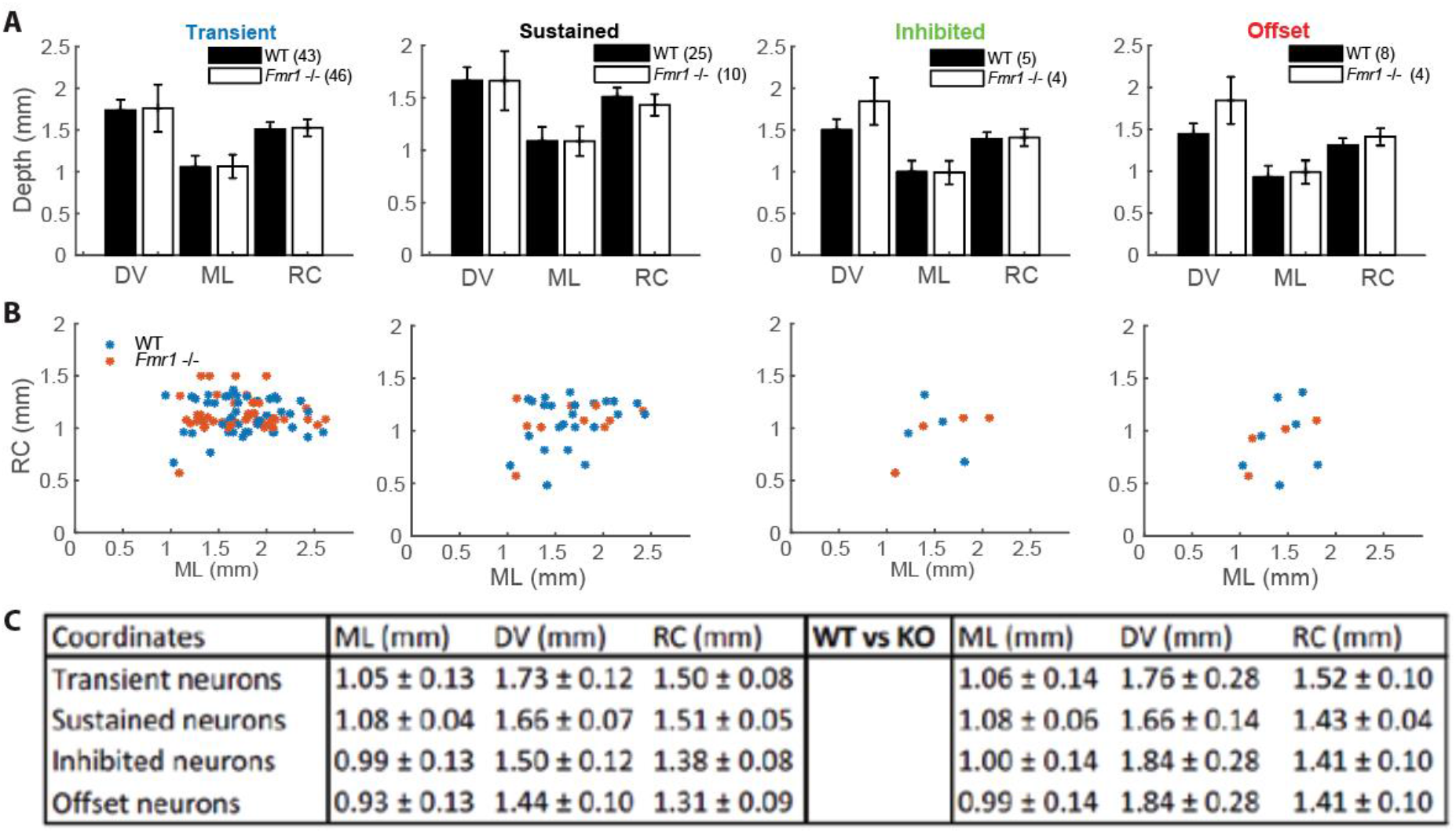
Detailed stereotactic recording locations in both genotypes. **A** Quantification of recording stereotactic depths in respectively: (left) standard neurons, (middle left) sustained neurons, (middle right) inhibited neurons, (right) offset neurons. **B** Quantifications of the respective Rostro-Caudal to Medio-Lateral stereotactic recording locations in resp.: (left) standard neurons, (middle left) sustained neurons, (middle right) inhibited neurons, (right) offset neurons. **C** Table of all values.

**Supplementary figure 2:**
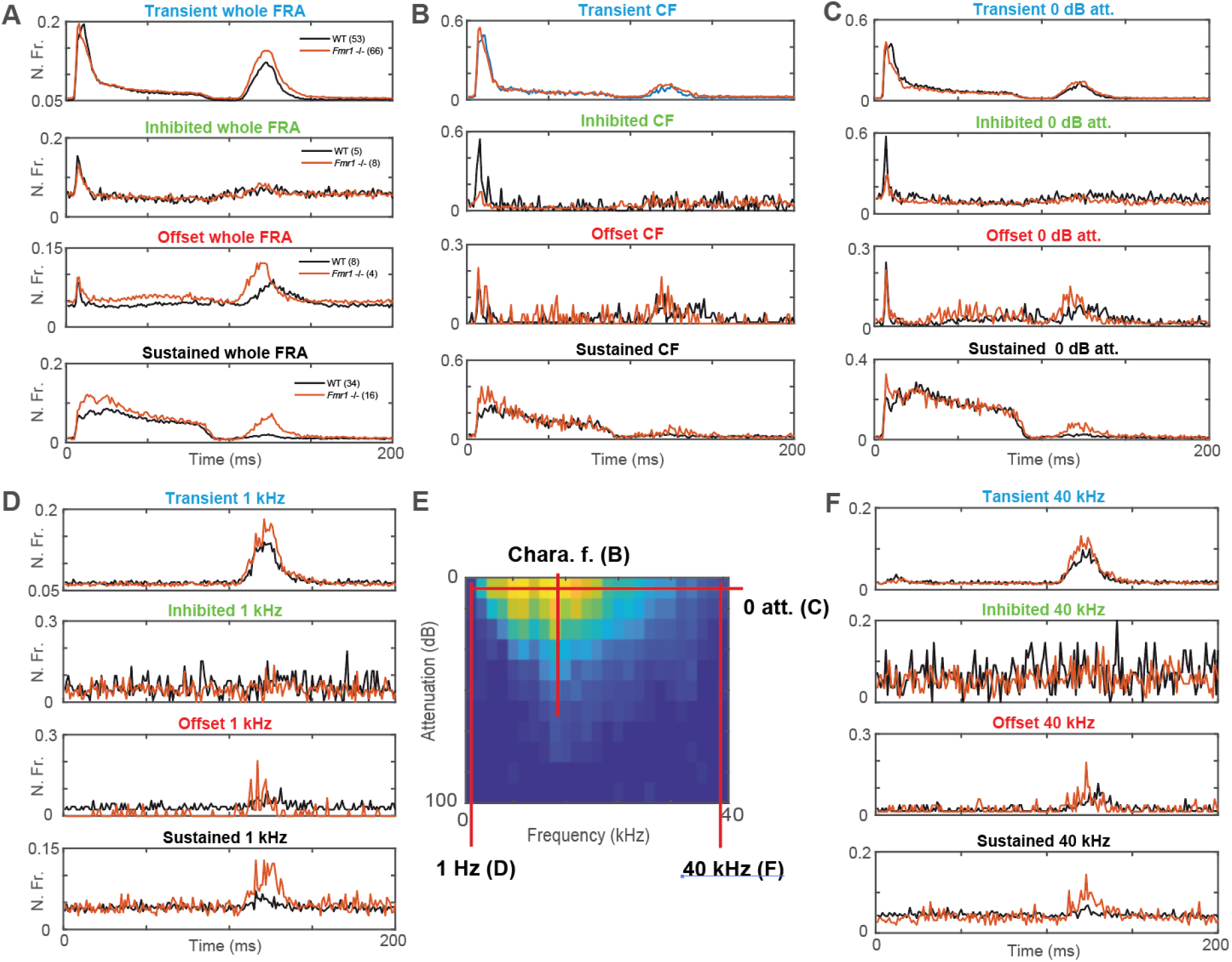
Localized estimates of the different firing patterns exhibited in defined sub- sections of the FRA. **A** Averaged compound PSTH of all conditions in the FRA for respectively: (above) all standard WT and all *Fmr1* -/- neurons, (middle above) all inhibited WT and all *Fmr1* -/- neurons, (middle below) all offset WT and all *Fmr1* -/- neurons, (below) all sustained WT and all *Fmr1* -/- neurons. **B** Averaged PSTH at the CF for all intensity below threshold for respectively: (above) standard WT and *Fmr1* -/-, (middle above) inhibited WT and *Fmr1* -/-, (middle below) offset WT and *Fmr1* -/-, (below) sustained WT and *Fmr1* -/-. **C** Averaged PSTH of 0 dB attenuation for all frequencies for respectively: (above) standard WT and *Fmr1* -/-, (middle above) inhibited WT and *Fmr1* -/-, (middle below) offset WT and *Fmr1* -/-, (below) sustained WT and *Fmr1* -/-. **D** Averaged PSTH of 1 kHz pure tone stimulations at all intensities for respectively: (above) standard WT and *Fmr1* -/-, (middle above) inhibited WT and *Fmr1* -/-, (middle below) offset WT and *Fmr1* -/-, (below) sustained WT and *Fmr1* -/-. **E**. Schematic overview of a FRA with red highlights the different sub-section chosen for the illustrated PSTH. **F** Averaged PSTH of 40 kHz pure tone stimulation for all intensities for respectively: (above) standard WT and *Fmr1* -/-, (middle above) inhibited WT and *Fmr1* -/-, (middle below) offset WT and *Fmr1* -/-, (below) sustained WT and *Fmr1* -/-.

**Supplementary figure 3:**
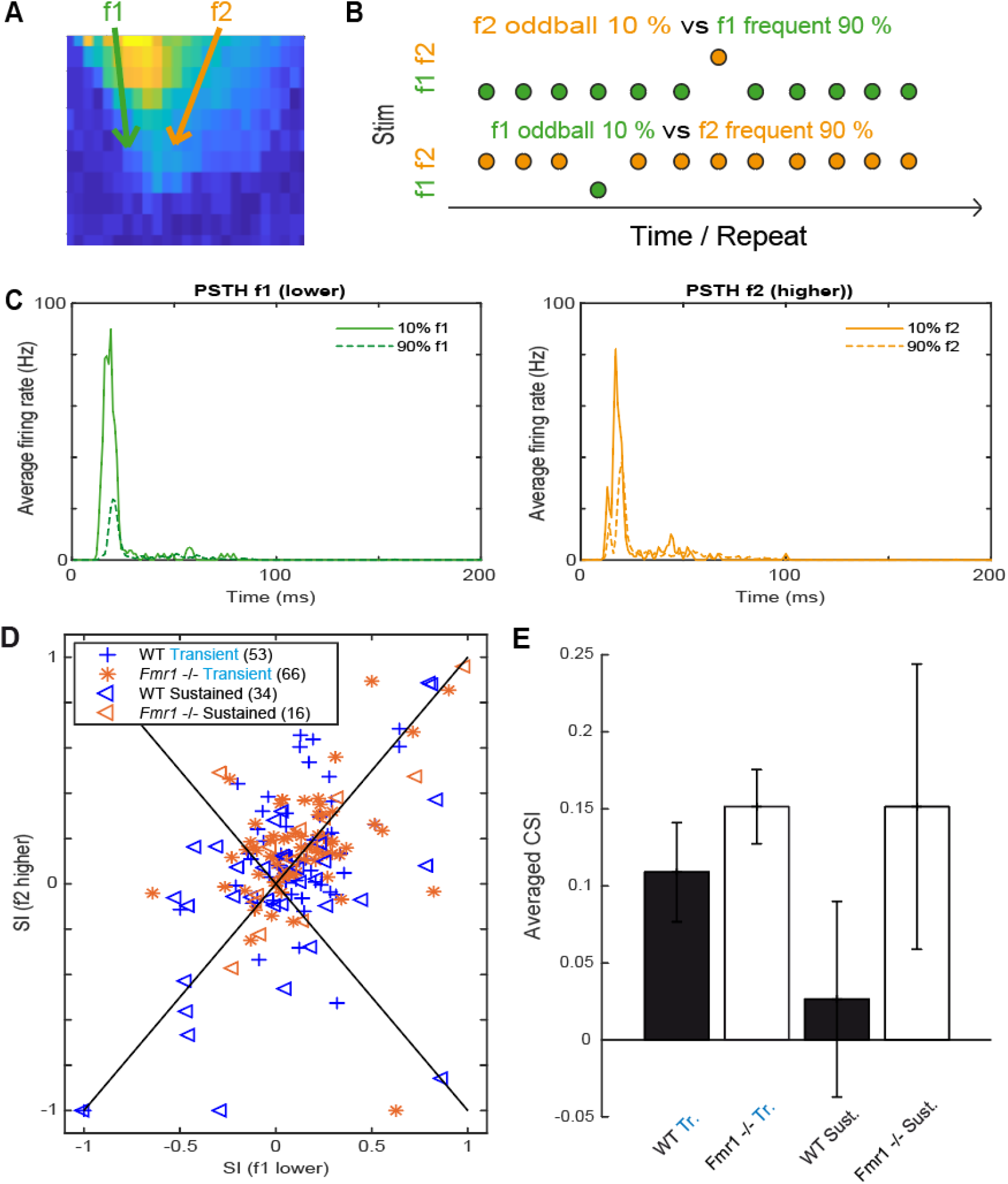
Neurons in the IC exhibit poor Stimulus Specific Adaptation. **A** Schematic FRA and the location around the CF chosen to identify f1 and f2. **B** Representation of (green and orange) the two frequencies of stimulation with the repeats used to bot oddball protocols. **C** PSTH to the different conditions in an adaptive neuron, (left) response to a (green) f1 odd, (dashed-green) f1 frequent stimulation; (right) responses to an (orange) f1 odd, (dashed-orange) f2 frequent stimulations. **D** Quantifications of the SSA to a standard 90/10 % oddball paradigm represented according to each neuron’s adaptive responses to both the lower frequency (f1-x axis) and the higher frequency (f2-y axis). **E** Averaged adaptation strength to both frequencies quantified by the Common SSA Index (CSI).

